# Integrated risk assessment reveals low biological activity of RNA-based crop protection formulations across plant and non-target systems

**DOI:** 10.64898/2026.06.18.732835

**Authors:** M. Dobrowitsch, S. Friedrich, J. Noetzold, M. Bender, K. Rybak, B. Moorlach, K. Eickelpasch, A. Patel, J. Wegener, A. C. U. Furch, S. Robatzek, C. Richter, B. M. Moerschbacher, A. Pich, A. Koch

## Abstract

RNA-based crop protection products are considered promising alternatives to conventional pesticides due to their high target specificity and selectivity profile. While current risk assessments mainly focus on sequence-dependent off-target effects, potential sequence-independent effects of RNA molecules and formulation components remain poorly understood. Here, we assessed the potential of naked and formulated RNA molecules to induce unintended biological responses in plants and non-target organisms using complementary plant stress and cellular toxicity assays. Single-stranded RNA (ssRNA), double-stranded RNA (dsRNA), circular RNA (circRNA), and chitosan-based RNA formulations were evaluated for their ability to induce early stress responses in plants and adverse effects in insect cells. Cytosolic Ca²⁺ signaling and reactive oxygen species (ROS) production were monitored in barley and Arabidopsis, while impedance-based phenotypic profiling (ECIS) was used to assess cellular responses in *Spodoptera frugiperda* (Sf21) cells. Neither naked nor formulated RNA preparations triggered pronounced Ca²⁺ influxes or ROS bursts, indicating an absence of substantial activation of early plant defense signaling. Likewise, RNA formulations caused only minor effects on insect cell attachment and spreading as a measure for cell viability. Observable responses occurred primarily at elevated concentrations of RNA formulations and were formulation-dependent rather than RNA-dependent. Overall, the investigated RNA molecules exhibited low biological activity across the tested non-target systems. Our results highlight the importance of evaluating RNA active ingredients and delivery matrices separately and demonstrate the value of integrated screening approaches for the early hazard assessment of RNA-based crop protection products.

## Introduction

The increasing restrictions on conventional chemical pesticides, together with the need for more sustainable crop protection strategies, have accelerated the development of novel, innovative plant protection products. Among these, RNA-based biopesticides have emerged as a promising technology due to their high target specificity and their potential to reduce unintended environmental impacts compared to broad-spectrum chemical compounds (Willow et al. 2025; Koch and Krczal 2026). RNA-based crop protection products typically exploit the natural mechanism of RNA interference (RNAi), in which double-stranded RNA (dsRNA) or related RNA molecules trigger sequence-specific degradation of target transcripts in pests or pathogens. Several RNA-based products are currently under development (Uslu et al. 2026), and the first RNAi-based insecticide (Yan et al. 2024) and acaricides (Narva et al. 2026) have already reached commercial application, demonstrating the feasibility of this approach in agriculture. As regulatory authorities worldwide establish frameworks for the environmental risk assessment of RNA-based products, a key challenge remains the evaluation of potential effects on non-target organisms (USEPA, 2023; OECD, 2023). Current risk assessments primarily focus on sequence-dependent off-target effects, assuming that biological activity is driven by complementarity between the applied RNA molecule and endogenous transcripts in exposed organisms (Petrick et al., 2013; De Shutter et al. 2022; Koch 2023; Koch and Krczal 2026; De Neef et al. 2025). Bioinformatic analyses have therefore become an important component of RNA pesticide safety evaluation (Uslu et al. 2026). However, considerably less attention has been paid to sequence-independent biological effects that may arise from the RNA molecule itself or from associated delivery systems. This knowledge gap is particularly relevant because nucleic acids can act as molecular patterns recognized by innate immune systems across a broad range of organisms, including plants, insects, and mammals (Maillard et al. 2013; Niehl et al., 2016; Huang et al. 2023; Hernández-Pelegrín et al. 2025).

In plants, extracellular nucleic acids have been reported to trigger defense-associated signaling pathways, including reactive oxygen species (ROS) production, cytosolic calcium influxes, and mitogen-activated protein kinase (MAPK) activation, which represent early hallmarks of pattern-triggered immunity (PTI) (Niehl et al., 2019; Heinlein 2025). Although such responses are generally weaker and less consistent than those induced by classical pathogen-associated molecular patterns, they raise the question whether externally applied RNA molecules could unintentionally activate stress or immune responses in non-target plant species. Similar concerns have been discussed for animal systems, where innate immune receptors are known to recognize foreign RNA molecules and initiate inflammatory signaling cascades (Kato et al., 2006; Shimizu 2024).

An additional aspect that can be overlooked in RNA-based product risk assessment is the contribution of formulation matrices. Due to the limited environmental stability and cellular uptake of naked RNA, various carriers and delivery systems are being developed to improve efficacy and persistence (Chen et al. 2023; Uslu et al. 2026). These formulations may include nanoparticles, lipids, polymers, or polysaccharides that possess biological activities independent of the RNA cargo. Chitosan-based formulations are among the most widely studied delivery systems because of their biodegradability, biocompatibility, and ability to protect nucleic acids from degradation (Kolge et al. 2021; Liu et al. 2024). However, chitosan is also recognized as a potent elicitor of plant defense responses and has been shown to induce ROS bursts, calcium signaling, stomatal closure, defense gene activation, and resistance-associated pathways in numerous plant species (El Hadrami et al., 2010; Melcher and Moerschbacher 2016; Gubaeva et al. 2018; Ye et al. 2020; Drs et al. 2025; Hellmann et al. 2026; Richter et al. 2025). Consequently, any safety assessment of RNA-formulations should distinguish between effects attributable to the RNA active ingredient and those originating from formulation additives. To address these knowledge gaps, we systematically evaluated potential sequence-independent biological responses induced by different RNA molecules, such as coding single stranded RNA (ssRNA), double stranded RNA (dsRNA) and circular RNA (circRNA) and RNA formulations in a range of relevant non-target test systems. Specifically, we investigated whether RNA compounds trigger early immune signaling responses in plants, including ROS production and cytosolic calcium influxes. Furthermore, we investigated potential biological responses in plant and insect-based test systems exposed to naked and formulated RNA preparations. Particular emphasis was placed on chitosan-containing formulations due to their increasing use in RNA delivery and their known immunomodulatory properties. The objectives of this study were therefore (i) to determine whether naked RNA molecules and RNA formulations induce sequence-independent stress or immune responses in plants, (ii) to distinguish between effects attributable to the RNA cargo and those originating from formulation matrices, and (iii) to evaluate the suitability of a combined assay battery comprising plant signaling and cell-based toxicity studies for the early risk assessment of RNA-based crop protection products.

## Material and Methods

### Synthesis of chitosan-adipic acid nanogels

Chitosan (95/20) (CTS, product line Chitoceuticals, degree of deacetylation ≥92.6 %, viscosity 16–30 mPas, molecular weight by gel permeation chromatography (GPC) 40–150 kDa) was obtained from Heppe Medical Chitosan GmbH. Adipic acid (AA) (99 %), Span 80 (sorbitan monooleate, mol wt 482.60 g mol⁻¹) and Tween 80 (polysorbate 80, mol wt 1310 g mol⁻¹) were purchased from Sigma Aldrich. Chitosan-adipic acid nanogels were synthesized by EDC-mediated crosslinking of chitosan (95/20) with adipic acid in an inverse mini-emulsion system. Nanogels with different crosslinking degrees (MB 22.1–26) were prepared by varying the ratio of the amino groups of chitosan to the carboxylic groups of adipic acid (composition of both phases is summarized in supplementary Table 1). Sonication was performed in pulse mode with alternating cycles of 3 s sonication and 1 s pause at an amplitude of 30 % for 2 min. The reaction mixture was stirred for 12 hat room temperature. The resulting nanogel suspension was purified, transferred into an aqueous solution, and subsequently characterized by dynamic light scattering (DLS), electrophoretic light scattering (ELS), and atomic force microscopy (AFM) (Supplementary Figs. S2–S3). The characterized formulations (MBX 22.1–26) were used for subsequent biological characterization and toxicity assessment.

### DLS and ELS Measurements

For dynamic light scattering (DLS) and electrophoretic light scattering (ELS) measurements, a ZetaSizer Ultra from Malvern Panalytics was usedand analysis was done with the ZS Xplorer software. To prepare the DLS samples, 100 µL of the nanogel suspension was transferred into a glass vial and diluted with 2 mL of HPLC grade water. The polystyrene cuvettes of the type DTS0012 were filled to approximately 1 cm with a syringe through a 1.2 µm PET syringe filter. ELS sample preparation was done identically and a cuvette of the type DTS1070 was used. All measurements were performed at 25 °C. Each sample was prepared independently in duplicate and each preparation was measured in triplicate. Each data point was averaged over these measurements.

### Scanning Force Microscopy Imaging

To prepare a sample for scanning force imaging (SFM), 100 µL of the nanogel suspension was transferred into a glass vial and diluted with 2 mL of HPLC grade water. A silicon wafer was cleaned with toluene and air dried prior to sample deposition. Afterwards, the wafer was treated with air-plasma for 180 s with a Flecto10USB-MFC plasma etcher by Plasma Technology to obtain a hydrophilic surface. A 50 µL droplet of nanogel suspension was spin-coated onto the wafer at 2000 rpm for 1 min on a WS-650SZ-&nPP/LITE by Laurell. Surface-coated nanogels were imaged with atomic force microscopy on a NanoScope V AFM by Veeco Instruments in tapping mode. The NCH-50 POINTPROBE-Silicon SPM-Sensor cantilever was manufactured by Nanoworld and had a resonance frequency of 320 kHz and a force constant of 42 N m-1. 10 µm x 10 µm images were evaluated in Gwyddion 2.70 to investigate particle morphology.

### Cytosolic Ca^2+^ measurements in barley leaf discs

Changes in cytosolic free Ca^2+^ concentration were monitored using transgenic *Hordeum vulgare* plants expressing cytosolic apoaequorin (Hv-AEQcyt #18, cv. Golden Promise), kindly provided by Edgar Peiter (Martin Luther University Halle-Wittenberg). The Hv-AEQcyt reporter line has previously been described for aequorin-based Ca^2+^ measurements in barley (Giridhar et al., 2022). Leaf discs were excised from barley leaves using a sterile razor blade and transferred individually into white 96-well microplates. Aequorin was reconstructedovernight in the dark by incubating leaf discs in 10 nM coelenterazine containing 0.01% (v/v) Tween-20. Luminescence measurements were performed using a Luminoskan Ascent luminometer (Thermo Electron Corporation, Finland). Baseline luminescence was recorded for 1 min prior to treatment. Subsequently, 50 µL of the respective test solution was added to each well and luminescence was recorded at 6-s intervals for 20 min. Distilled water served as the negative control, whereas chitohexaose was used as a positive control for elicitor-induced Ca^2+^ signaling. Following each measurement, residual aequorin was discharged by addition of 1 M CaCl_2_ in 10% (v/v) ethanol. Cytosolic Ca^2+^ concentrations were calculated from relative luminescence values as described by Mithöfer and Mazars (2002). For comparative analyses, peak Ca^2+^ responses were normalized to the mean chitohexaose response obtained within each experiment.

### ROS assays in leaf discs

ROS production was measured in white non-transparent 96-well plates (Greiner Bio-One; 665075) in an ICCD photon counting camera (Photek, East Sussex, UK). Two 96-well plates were measured at the same time. Leaf discs (diameter 4 mm) from 2-week-old barley (*Hordeum vulgare* cv. Golden Promise; cultivated at 16 hours light period with 18°C during the day and 14°C during the night, humidity was set at 65% and light intensity was 240 µmol m⁻² s⁻¹with a ratio of warm white light to cool white light of 1:1) or 4-week-old *Arabidopsis thaliana* plants were harvested one day before the assay. Per treatment, eight technical replicates were conducted, consisting of leaf discs from at least three individual plants. Leaf discs were placed in wells containing 150 µl distilled water (dH_2_O) and incubated with transparent lids closed overnight at room temperature. The next day, the water was removed with a multi-tips pipette. For *A. thaliana*, the assay solution consisted of 17 µg/ml Luminol (17 mg/ml stock in DMSO; A8511; Sigma) and 10 µg/ml horseradish peroxidase (10 mg/ml stock in water; P6782; Sigma). For barley, the assay solution consisted of 34 µg/ml Luminol L-012 (17 mg/ml stock in water; FUJIFILM Wako Chemicals; 120-04891) and 10 µg/ml horseradish peroxidase (10 mg/ml stock in water; 77332; Sigma). For *A. thaliana*, 1 µM flg22 (1 mM stock; Peptron) was used as positive control. For barley, 0.2 mg/ml chitin (10 mg/ml stock in water; C9752; Sigma) was used as positive control. Additionally, 50 µg/ml Chitosan (2 mg/ml stock, dissolved in dH_2_O containing 5% molar excess of acetic acid relative to the free amino groups; DA 20, DP 500; Gillet Chitosan SAS, France) was tested as a potential positive control but did not trigger a ROS burst in *A. thaliana* or barley. One hundred µl of assay solution was added to each well and the measurement was started immediately. Each assay was conducted for 2 h. For display of relative light units (RLU) one data point per 60 seconds was chosen, culminating in 120 data points per well. For determination of maximum RLU value (RLU_max_) of each well, the first 10 data points were ignored, since the mechanic trigger through the addition of the assay solution caused a non-specific ROS response. For each treatment, consisting of eight technical replicates, the mean and standard deviation were calculated.

### Impedance-based Phenotypic Assay

To assess the phenotypic impact of RNA-based pesticides and microgel formulations on non-target organisms, Sf21 insect cells (provided by the Department of Biochemistry III of the University of Regensburg from the lab of Prof. Dr. Gernot Laengst) were monitored by electric cell-substrate impedance sensing (ECIS) during exposure. The principle of the assay has been described in detail previously (Friedrich et al., 2024). It provides a holistic, dose-dependent response of the cells upon exposure to the substances under test. In contrast to target-specific assays, the ECIS readouts integrate over the entire cell body and mirror the cells’ overall physiological response. The ECIS-assay is based on seeding cells on thin gold-film electrodes while applying a non-invasive, weak alternating current. During cell spreading, adherent cells impede the current due to the dielectric properties of their cell membranes, resulting in time-resolved changes in the complex AC impedance. The imaginary part of the impedance (reactance) at a sampling frequency of 20 kHz is expressed as the equivalent capacitance. The capacitance at this frequency scales linearly with electrode coverage (Wegener et al., 2000). The electrical readout is holistic as it reads complex phenotypic responses such as cell adhesion, morphology, or viability. In terms of interpretation low capacitance values correspond to electrodes confluently covered with tightly adherent and physiologically intact cells indicating no or little adverse effects of the test substances. In contrast, high capacitance values indicate cell-free or sparsely covered electrodes, reflecting cytotoxicity of the test compound and the resulting loss of cell adhesion and viability (Supplementary Figure 1). To investigate the off-target effects of RNA-based pesticides and microgel formulations, Sf21 insect cells (derived from *Spodoptera frugiperda*) were exposed to the various samples in dose-response studies using ECIS as a time-resolved readout (cell culture conditions are summarized supplemental Table 2). Sf21 cells were routinely grown in suspension, and subcultivated by diluting cells into fresh Erlenmeyer flasks. To assess toxicity of test substances, cell suspensions were centrifuged, resuspended in fresh medium and diluted to intended seeding density providing a confluent monolayer after attachment (300,000 cells/cm²). For the ECIS-based phenotypic assays, we used electrode arrays purchased from Applied BioPhysics Inc. (Troy, NY) of type 96W10idf. First electrode arrays were sterilized using argon plasma treatment before the baseline impedance of a cell-free electrode was recorded. Cells were then seeded onto the electrodes while being simultaneously mixed with the respective test substances, ensuring continuous exposure from the onset of adhesion and attachment. Cell attachment and spreading was identified as the most sensitive cell phenotype to assess adverse effects. Samples were diluted in culture medium using an aqueous stock solution. Data are presented as bar graphs showing area under the curve values (AUC) obtained from integration of capacitance time courses (0 to 5 h) at a frequency of 20 kHz. AUC values were corrected for the corresponding AUC of cell-free electrodes and normalized to AUC of untreated cells. Further methodological details are provided by Friedrich et al. (2024).

## Results

### Characterization of chitosan–adipic acid nanogels

Prior to biological testing, the physicochemical properties of the chitosan–adipic acid nanogels were characterized by dynamic light scattering (DLS), electrophoretic light scattering (ELS), and atomic force microscopy (AFM) (Supplementary Figs. S2–S3). The synthesized nanogels exhibited hydrodynamic diameters of approximately 300–500 nm and negative zeta potentials after purification, consistent with stable aqueous dispersions. Variation of the crosslinking density and surfactant concentration affected particle size distribution but did not substantially alter the overall colloidal properties of the formulations. These characterized nanogel formulations were subsequently used for biological testing.

### RNA formulations do not trigger early calcium-mediated stress responses

To assess whether RNA-based plant protection formulations induce early cellular stress responses, cytosolic Ca²⁺ transients were monitored following exposure to formulated and non-formulated RNA preparations. In all experiments, chitohexaose elicited the expected strong calcium response and served as a functional positive control, whereas dH₂O treatment resulted in low basal Ca²⁺ levels (Fig. 1A,B; Fig. 2A). Across all tested MBX formulations (MBX 22.1–26), calcium peak amplitudes remained similar to the dH₂O control and substantially below the response induced by chitohexaose (Fig. 1A). Although minor variation between formulations and concentrations was observed, no consistent concentration-dependent increase in cytosolic Ca²⁺ levels was detected. Similarly, CA-based carrier formulations loaded with short RNA, long RNA, or dsRNA did not induce elevated calcium responses relative to empty carrier controls (Fig. 1B). Calcium peak amplitudes remained within the range of the negative control irrespective of RNA cargo or concentration, indicating that neither the carrier material nor RNA loading triggered detectable early calcium signaling. Exposure to naked RNA molecules, ranging from 0,1 to 10 µg, likewise failed to induce pronounced cytosolic Ca²⁺ elevations (Fig. 2A). Both ssRNA and dsRNA preparations produced calcium peak amplitudes comparable to the water control across the tested concentration range. A comparative overview of all treatments further demonstrated that calcium responses remained clustered near the negative control and were consistently separated from the strong response elicited by the positive control (Fig. 2B).

**Figure 1.**
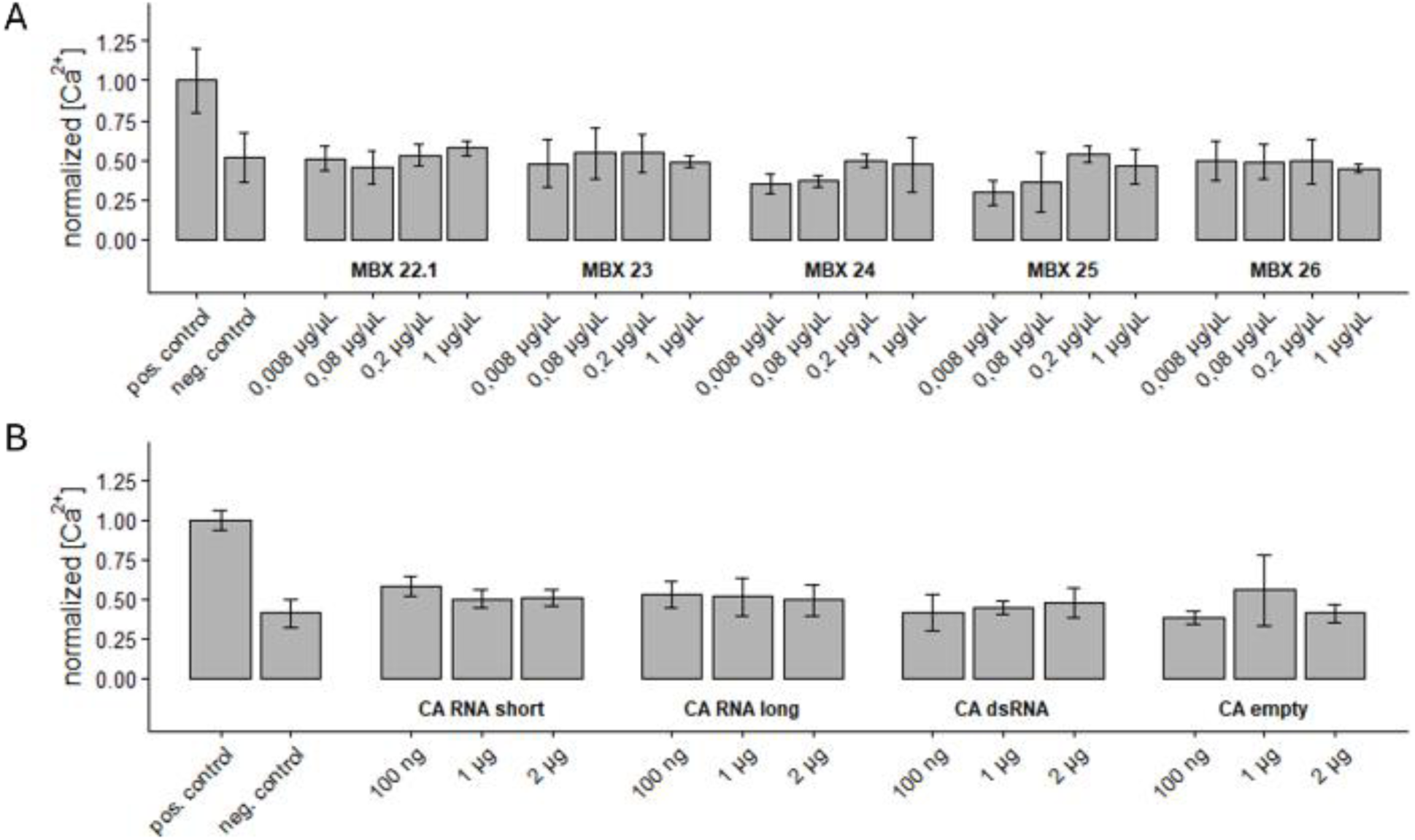
RNA formulations do not induce elevated cytosolic Ca²⁺ responses in barley leaf tissue. **(A)** Normalized peak cytosolic Ca²⁺ responses in transgenic aequorin-expressing barley leaf discs following treatment with chitosan-based nanogel formulations MBX 22.1–MBX 26 at the indicated concentrations. Chitohexaose served as a positive control and dH₂O as a negative control. **(B)** Normalized peak cytosolic Ca²⁺ responses following treatment with chitosan–alginate (CA) formulations containing short RNA, long RNA, or dsRNA, as well as empty carrier controls, at the indicated RNA concentrations. Chitohexaose and dH₂O served as positive and negative controls, respectively. Peak Ca²⁺ amplitudes were normalized to the mean response elicited by chitohexaose within each experiment. Bars represent mean ± SD of n = three biological replicates.

**Figure 2.**
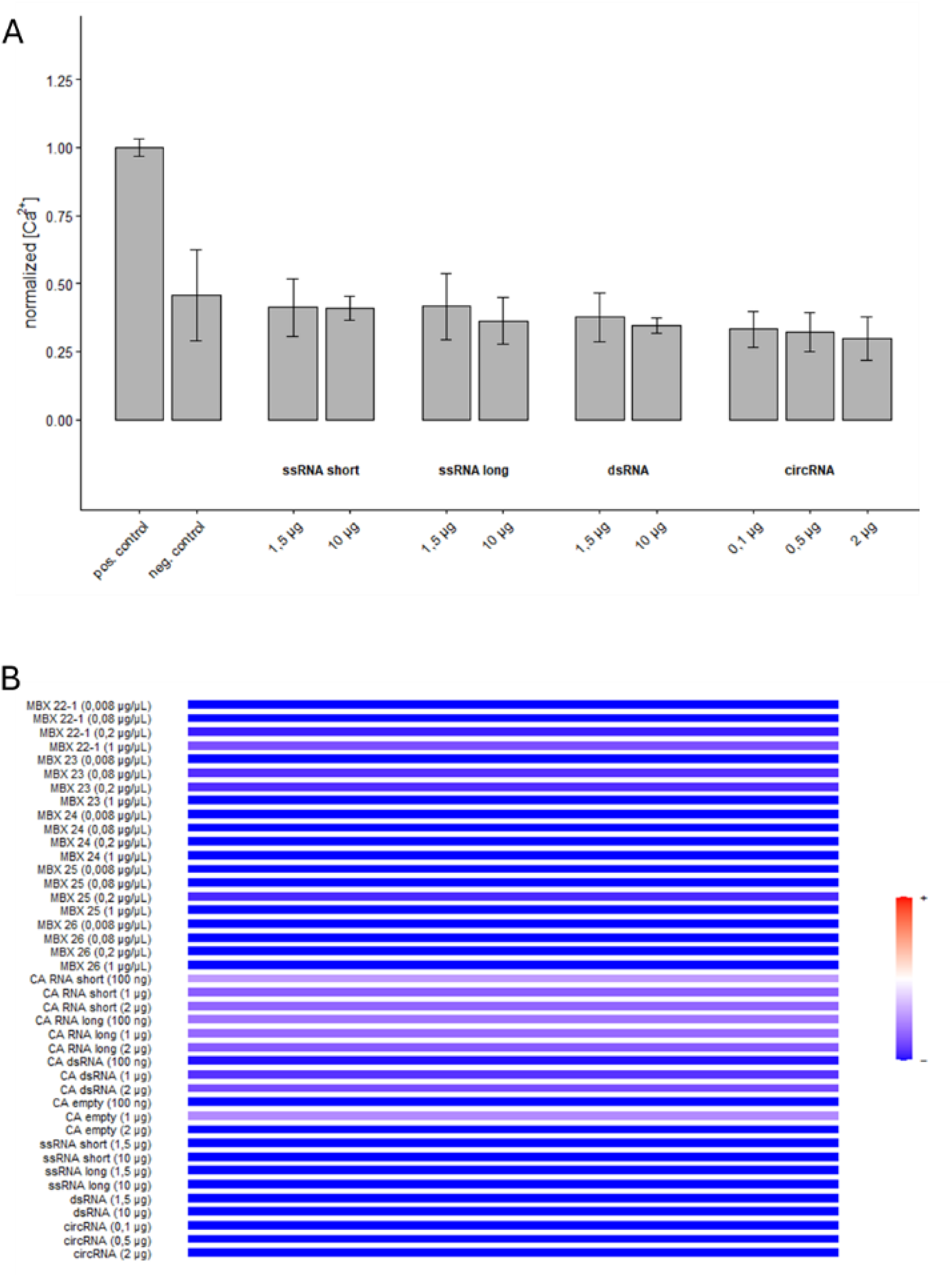
Cytosolic calcium responses following exposure to naked RNA molecules and summary of all tested RNA formulations. **(A)** Normalized peak cytosolic Ca²⁺ responses in transgenic aequorin-expressing barley leaf discs following treatment with naked RNA molecules. Responses were determined for short single-stranded RNA (ssRNA short), long single-stranded RNA (ssRNA long), double-stranded RNA (dsRNA), and circular RNA (circRNA) at the indicated concentrations. Chitohexaose and dH₂O served as positive and negative controls, respectively. **(B)** Heatmap summarizing normalized peak cytosolic Ca²⁺ responses across all tested RNA formulations, carrier systems, and naked RNA preparations. Color intensity reflects the relative magnitude of the normalized Ca²⁺ response, with blue indicating low responses and warmer colors indicating higher responses. All treatments remained clearly separated from the positive control and clustered near the negative control. Peak Ca²⁺ amplitudes were normalized to the mean response elicited by chitohexaose within each experiment. Bars represent mean ± SD of n = three biological replicates.

Collectively, these results indicate that neither the investigated RNA molecules nor their formulation platforms activate substantial calcium-dependent stress or defense signaling under the conditions tested.

### Limited ROS responses following exposure to chitosan-based RNA formulations

Reactive oxygen species (ROS) production was monitored to evaluate whether chitosan-based RNA delivery systems trigger early stress or defense responses in plants. In all experiments, treatment with the bacterial elicitor flg22 induced a strong ROS burst and served as a functional positive control, whereas mock-treated samples exhibited only low basal ROS levels (Fig. 3A, B). For *Arabidopsis*, across all tested chitosan microgel formulations (MBX 22.1–26), ROS production remained close to mock levels and substantially below the response induced by flg22 (Fig. 3A). No consistent concentration-dependent increase in ROS accumulation was observed over the tested concentration range. Although minor variation between individual formulations was detected, none of the formulations elicited ROS responses indicative of strong activation of plant defense signaling.

**Figure 3.**
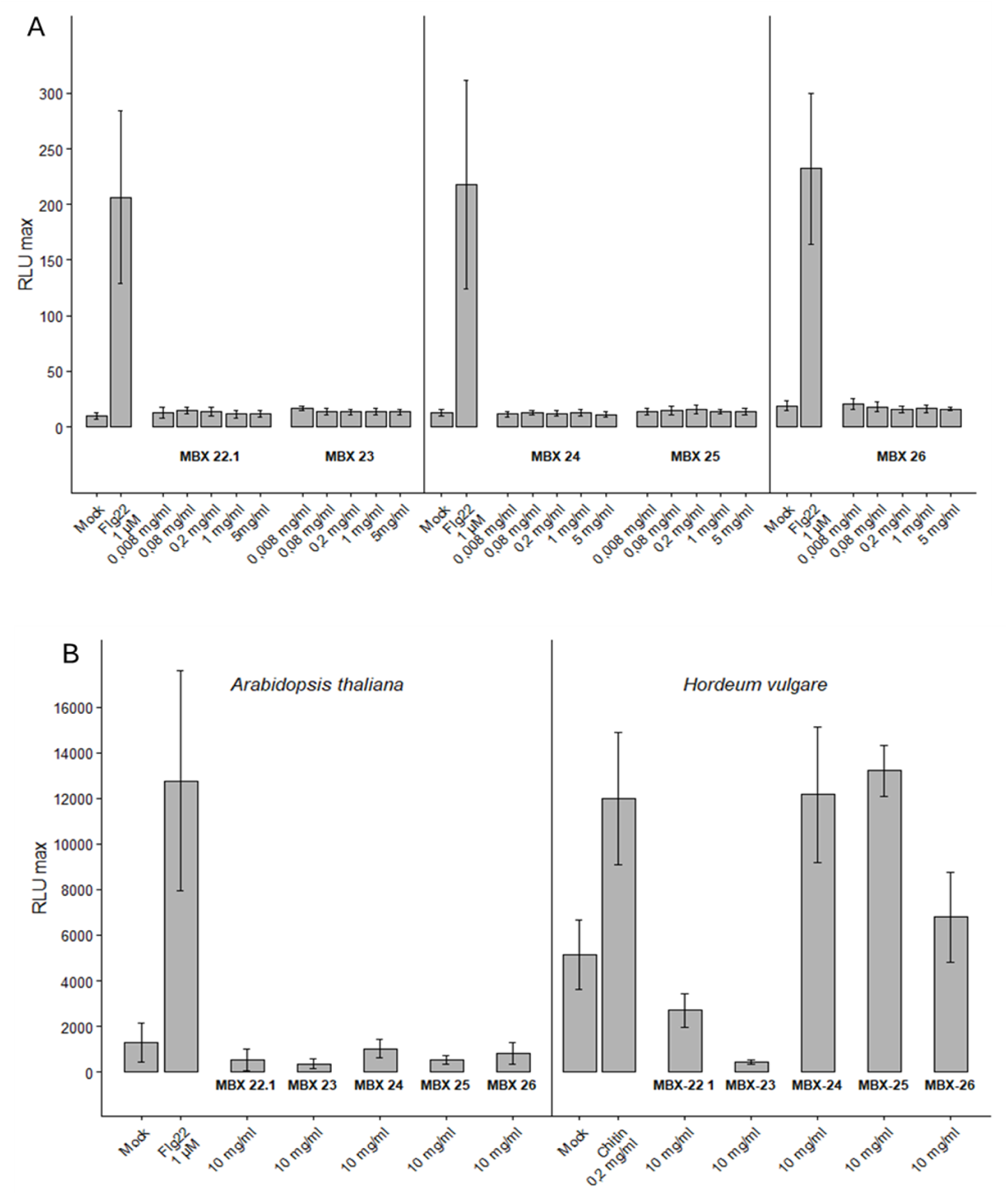
ROS responses to chitosan-based RNA formulations in Arabidopsis and barley. **(A)** Reactive oxygen species (ROS) production in *Arabidopsis thaliana* leaf discs following exposure to chitosan–microgel formulations MBX22.1–MBX26 at the indicated concentrations. ROS accumulation is shown as maximum relative luminescence units (RLUmax) measured using a luminol-based assay. Mock-treated samples served as negative controls, whereas flg22 (1 µM) served as a positive control for elicitor-induced ROS production. None of the tested formulations induced ROS levels comparable to the positive control, and no consistent concentration-dependent increase in ROS production was observed. **(B)** ROS production in *Arabidopsis thaliana* and *Hordeum vulgare* following treatment with the highest formulation concentration (10 mg mL⁻¹). Mock-treated samples served as negative controls. Flg22 (1 µM) and chitin (0.2 mg mL⁻¹) were used as positive controls for *Arabidopsis* and barley, respectively. Data are presented as mean ± SD of *n* = three biological replicates.

To assess whether potential responses were species-specific, the highest formulation concentration (10 mg mL⁻¹) was additionally evaluated in both *Arabidopsis* and barley (Fig. 3B). Consistent with the concentration series, all formulations induced only weak ROS responses in *Arabidopsis*. In contrast, formulation-dependent differences became apparent in barley. While MBX-23 remained close to basal levels, MBX-22.1 induced a moderate ROS response, and MBX-24 and MBX-25 triggered substantially elevated ROS production approaching the level of the chitin positive control. MBX-26 produced an intermediate response. These findings indicate that ROS induction by the investigated formulations is species- and formulation-dependent.

Taken together, the results demonstrate that the tested chitosan-based formulations do not induce pronounced ROS responses in *Arabidopsis*, whereas selected formulations can trigger measurable oxidative burst responses in barley at high concentrations. The absence of strong ROS induction in most treatments, together with the lack of elevated Ca²⁺ signalling, suggests a generally low potential for activation of early plant stress responses, although formulation-specific effects cannot be excluded.

### ECIS-based phenotypic profiling indicates low toxicity of RNA formulations in Sf21 insect cells

Potential adverse effects of RNA-based pesticides and formulation components were assessed in Sf21 cells derived from *Spodoptera frugiperda* using impedance-based phenotypic profiling. Cellular responses were quantified as normalized area-under-the-curve (AUC) values derived from capacitance time courses, with reduced AUC values indicating impaired cell attachment, spreading, viability, or morphology. Exposure to naked RNA preparations resulted in only minor effects across the tested concentration range (Fig. 4A). For dsRNA, short RNA and long RNA, normalized AUC values remained close to those of untreated controls at low and intermediate concentrations. Moderate reductions were observed only at the highest tested RNA concentrations, while values remained clearly above those of cell-free controls, indicating limited adverse effects on cellular integrity. Similarly, formulated RNA preparations induced only moderate phenotypic effects in Sf21 cells (Fig. 4B). At the lower tested concentration, normalized AUC values remained similar to untreated controls irrespective of RNA type. At the higher concentration, reductions in AUC values were observed for all formulated RNA preparations, indicating impaired cell attachment and spreading. Because similar responses were observed for the corresponding RNA-free formulation control, these effects are likely attributable primarily to the formulation matrix rather than the RNA cargo itself.

**Figure 4.**
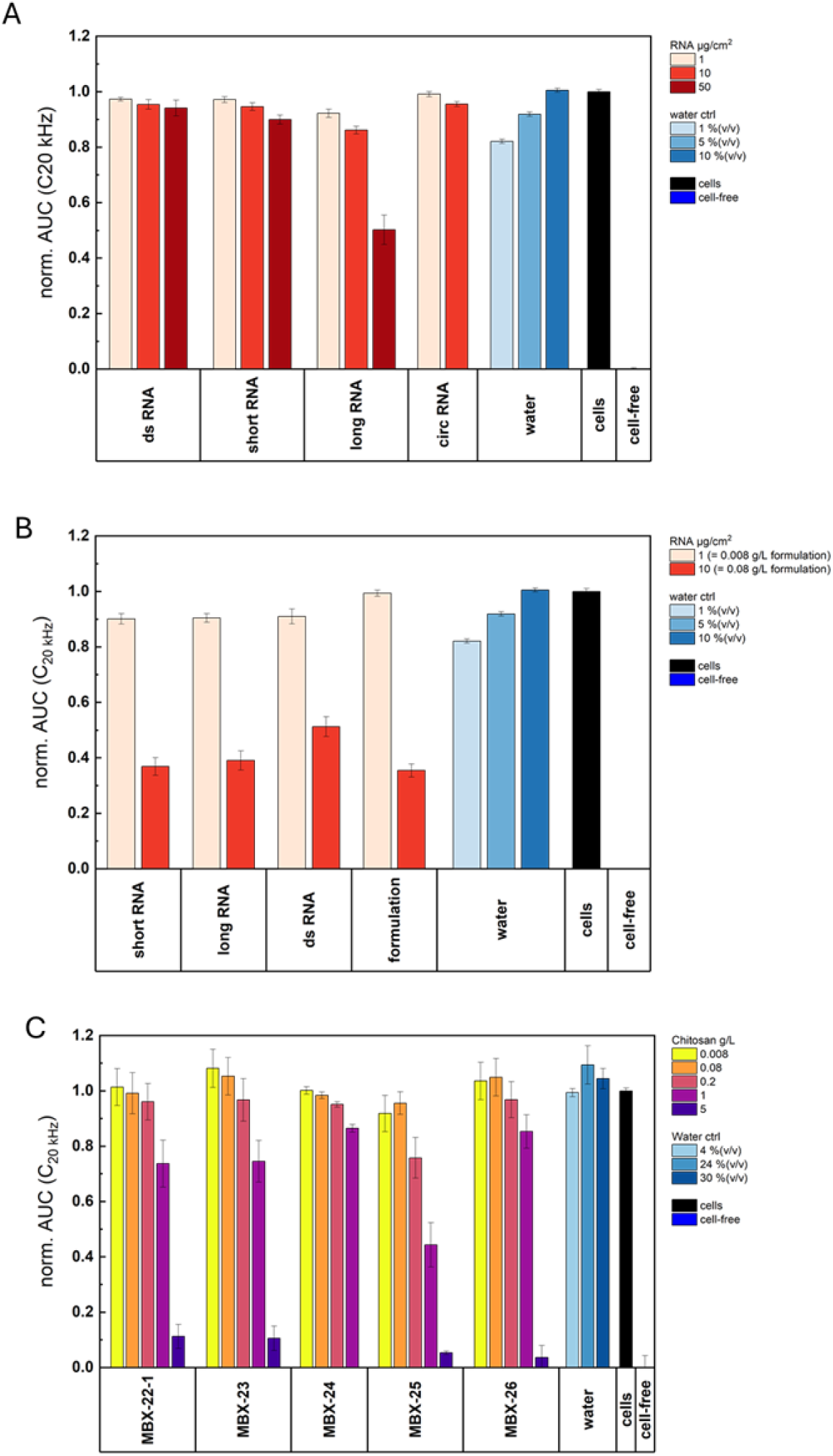
ECIS-based phenotypic profiling of RNA formulations in Sf21 insect cells. **(A)** Effects of naked RNA molecules on Sf21 cell attachment and spreading. Cells were exposed to short single-stranded RNA (short RNA), long single-stranded RNA (long RNA), double-stranded RNA (dsRNA), or circular RNA (circRNA) at the indicated surface concentrations. Cellular responses are shown as normalized area-under-the-curve (AUC) values derived from capacitance measurements at 20 kHz. **(B)** Comparison of formulated RNA preparations and formulation controls. Sf21 cells were exposed to chitosan–alginate microgel formulations containing RNA cargo or corresponding empty formulations at concentrations equivalent to the indicated RNA loading. Formulation controls were tested at concentrations corresponding to the respective RNA-loaded formulations. **(C)** Effects of empty chitosan–nanogel formulations (MBX 22.1–MBX 26) on Sf21 cells. Cells were exposed to increasing concentrations of the respective formulation materials in the absence of RNA cargo. Water controls correspond to the dilution range introduced by the tested formulations. Data are presented as normalized AUC values relative to untreated cell controls. Reduced AUC values indicate impaired cell attachment, spreading, viability, or morphology. Bars represent mean ± SEM of n = 4 to 7 (one experiment. Cell-free electrodes served as the assay reference for complete loss of cell coverage.

In line with this, exposure to formulation material in the absence of RNA revealed concentration-dependent effects for the chitosan–nanogels (Fig. 4C). While low and intermediate concentrations produced responses similar to untreated controls, higher concentrations resulted in reduced normalized AUC values, indicating impaired cell attachment and spreading. The magnitude of these effects varied between formulations but remained less pronounced than the readings for cell-free electrodes.

Overall, the ECIS-based phenotypic analysis indicates low intrinsic toxicity of the tested RNA molecules towards Sf21 cells. Phenotypic effects became apparent primarily at elevated concentrations and were largely associated with the formulation material rather than the RNA cargo itself.

## Discussion

Calcium influx and ROS production are among the earliest cellular responses following perception of pathogen-associated or damage-associated molecular patterns, including bacterial elicitors such as flg22 and fungal cell wall-derived molecules such as chitin or chitosan fragments. The absence of elevated Ca²⁺ transients across all tested treatments, together with the generally low ROS responses observed in Arabidopsis and the limited, formulation-dependent ROS responses detected in barley, indicates that the investigated RNA formulations were not perceived as strong activators of early plant defense signaling under the conditions applied. This finding is relevant because exogenous dsRNA has previously been proposed to act as a pathogen-associated molecular pattern in plants. Niehl et al. (2016) reported that dsRNA can induce a PTI-like signaling pathway in *Arabidopsis* involving SERK1 and a putative dsRNA receptor, independent of canonical Dicer-like RNA silencing pathways. In contrast, our data do not support a generalized activation of early PTI-like responses by externally applied RNA. Neither naked dsRNA nor ssRNA induced Ca²⁺ or ROS responses comparable to the positive controls, and formulation of RNA did not enhance these responses (Fig 1-3). These apparently divergent observations may reflect differences in RNA structure, length, concentration, purity, mode of application, plant species, tissue type, and formulation context. In particular, formulation may alter the accessibility of RNA molecules to putative pattern-recognition systems. Moreover, current reviews on exogenous RNA applications emphasize that uptake, stability, processing, and biological activity of externally applied dsRNA are highly context-dependent rather than uniform across systems (Uslu et al. 2026). Thus, our results do not exclude that specific dsRNA molecules or application conditions may activate immune signaling, but they indicate that such activation is not an inherent property of all externally applied RNA preparations. Previous studies further suggest that activation of plant immunity by exogenous RNA is highly context-dependent. While Niehl et al. (2016) and more recently Huang et al. (2023) provided evidence that specific dsRNA molecules can trigger PTI-like responses, including defense-associated signaling and regulation of cell-to-cell transport, such responses do not appear to be universally associated with exogenous RNA applications. Recent findings by Zheng et al. (2025) further support this view by demonstrating that biological responses to externally applied dsRNA depend strongly on dose, target organism, and formulation context. In the fungus *Magnaporthe oryzae*, high dsRNA concentrations induced sequence-independent stress responses, including ROS production and activation of the Hog1p MAPK pathway, whereas sequence-specific RNAi effects predominated at lower concentrations. Moreover, alginate–chitosan nanoparticles enhanced the protective efficacy of dsRNA, highlighting the importance of formulation-dependent biological activity. Together, these findings indicate that exogenous dsRNA can elicit both sequence-specific and sequence-independent responses, but that the occurrence and biological relevance of such effects vary considerably among biological systems.

Evidence from SIGS studies likewise suggests that effective RNA-based disease control does not necessarily require activation of canonical plant defense pathways. In a pioneering SIGS study, Koch et al. (2016) investigated whether exogenously applied CYP3-dsRNA induces defense-associated responses in barley. Despite conferring protection against *Fusarium graminearum*, CYP3-dsRNA did not induce expression of the salicylic acid-responsive marker gene *HvPR1* or the jasmonate-responsive marker *HvJMT*, whereas both genes were strongly induced following fungal infection. These findings led the authors to conclude that SIGS-mediated disease protection operates independently of canonical innate immune activation. The absence of elevated Ca²⁺ transients and ROS bursts observed in the present study is consistent with this interpretation and further supports the view that exogenous RNA can exert biological activity without necessarily triggering a detectable PTI response in the host plant. Collectively, the available evidence suggests that dsRNA perception and immune activation are not inherent consequences of exogenous RNA application, but rather depend on multiple factors including RNA structure, concentration, formulation, target organism, and exposure conditions.

The formulation component is also important in this context. Chitosan and chitosan-derived oligomers are widely described as plant defense elicitors capable of inducing ROS production, MAPK signaling, defense-related gene expression, and immune priming, depending on their physicochemical properties and application conditions. Recent work further suggests that receptor-mediated bioactivity of chitosans is influenced not only by the overall degree of acetylation but also by the underlying pattern of acetylation and the production process itself. Hellmann et al. (2026) demonstrated that chitosans produced by different chemical processing routes (heterogeneous deacetylation of chitin versus chemical N-acetylation of polyglucosamine) can exhibit substantially different receptor-mediated bioactivities despite similar bulk physicochemical properties, highlighting that biological activity depends not only on overall composition but also on structural features such as the pattern of acetylation. These findings indicate that biological activity cannot be predicted solely from conventional descriptors such as molecular weight or degree of deacetylation and suggest that subtle structural features of chitosan polymers may critically influence recognition by chitin- and chitosan-responsive receptors.

Against this background, the absence of pronounced Ca²⁺ and ROS responses following application of the chitosan-containing formulations is notable. Rather than indicating that chitosan is intrinsically inactive, our data suggest that some specific chitosan–architecture strongly attenuates the elicitor activity commonly associated with soluble chitosan preparations. Complexation with alginate, reduced polymer accessibility, altered surface charge characteristics, particle size effects, or limited release of bioactive chitosan fragments may all contribute to reduced recognition by plant perception systems. The observed lack of activity may therefore not only result from encapsulation within the alginate matrix but also from intrinsic structural properties of the chitosan used for particle preparation. Together, these observations support the emerging view that chitosan-containing delivery systems cannot be considered biologically equivalent and should be evaluated on a formulation-specific basis.

From a risk-assessment perspective, these findings highlight the importance of evaluating RNA active ingredients and formulation components separately. Chitosan-containing carriers may combine desirable formulation properties, such as improved RNA protection and delivery, with the potential to modulate plant physiology. Consequently, safety assessments of RNA-based plant protection products should address both the RNA cargo and the delivery system. Under the tested conditions, neither empty nor RNA-loaded formulations induced strong or consistent early immune-like responses. While no substantial Ca²⁺ responses were detected and ROS production remained low in Arabidopsis, some formulation-dependent ROS responses were observed in barley. Overall, these findings indicate a low potential of the investigated formulations to act as acute elicitors of early plant defense signaling. Nevertheless, the absence of pronounced early responses does not exclude more subtle or delayed physiological effects. Future studies may therefore benefit from targeted analyses of defense-associated marker genes or priming responses at later time points following treatment.

While early plant signaling assays provide information on potential plant stress responses, the ECIS-based phenotypic profiling approach offers a complementary perspective by assessing direct effects on non-target cells. Unlike endpoint-specific toxicity assays, impedance-based measurements integrate multiple cellular properties, including attachment, spreading, morphology, and viability, thereby providing a holistic measure of cellular fitness. Across the tested concentration range, naked RNA preparations induced only minor effects on cellular behavior (Fig 4). Normalized AUC values remained close to untreated controls at low and intermediate concentrations, and only moderate reductions were observed at the highest exposure levels (Fig 4C). Importantly, these responses remained clearly separated from the cell-free control, indicating that cellular integrity was largely maintained even under worst-case exposure scenarios. These observations are consistent with the generally accepted view that RNA molecules themselves possess limited intrinsic toxicity because they are naturally occurring biomolecules that are rapidly degraded in biological and environmental systems (OECD 2023). A particularly relevant finding is that formulation of RNA within chitosan–alginate does affect adverse cellular responses compared to the corresponding naked RNA preparations (Fig. 4). Across all tested RNA types, the effect was significant only for the higher concentrations under test. For the lower concentrations formulated and non-formulated RNA were highly similar, suggesting that encapsulation did not introduce additional toxicity towards the investigated non-target cell models at low concentrations but became relevant at higher doses. This observation is important because regulatory concerns associated with RNA-based pesticides increasingly focus not only on the active RNA ingredient but also on delivery technologies that may alter exposure, uptake, persistence, or biological interactions (De Neef et al. 2025; Uslu et al. 2026; Koch and Krczal 2026). In contrast, the strongest responses observed in this study were associated with the formulation matrix itself rather than the RNA cargo (Fig 4). Although these effects were generally limited and became apparent only at elevated concentrations, they indicate that formulation components can contribute more substantially to biological responses than the encapsulated RNA molecules. Comparison between formulated RNA and formulation only indicates that the formulation matrix is responsible for the phenotypic response.

Taken together, the ECIS data indicate that the tested RNA molecules exhibit low intrinsic toxicity towards Sf21 insect cells and that observed effects are primarily associated with high concentrations of formulation material rather than the RNA cargo itself. These findings further support the importance of evaluating RNA active ingredients and formulation matrices separately when assessing the biological activity of RNA-based crop protection products. The ECIS assay therefore provides a useful tool to discriminate between RNA- and formulation-associated effects during early-stage safety evaluation. This is particularly relevant as ECIS assays are scalable to high-throughput formats up to 600 samples in parallel.

The regulatory evaluation of RNA-based plant protection products is currently undergoing rapid development, with increasing emphasis on both RNA active ingredients and their associated delivery systems (OECD, 2020; OECD, 2023; De Schutter et al., 2022; Koch and Krczal 2026). While substantial progress has been made in the assessment of sequence homology-based off-target risks and environmental exposure scenarios (De Neef et al. 2025), comparatively little attention has been given to experimental approaches that address sequence-independent biological activity and formulation-related effects during early product development (Uslu et al. 2026). The present study demonstrates that early biological screening can provide valuable information on potential unintended effects of RNA-based plant protection products beyond sequence-specific off-target considerations. Across three independent biological endpoints, including plant calcium signaling, ROS production, and ECIS-based phenotypic profiling, the investigated RNA formulations consistently exhibited low biological activity towards the tested plant and cell-based systems. A key finding of this study is that biological responses were more strongly associated with formulation characteristics than with the RNA molecules themselves. While naked RNA preparations showed little evidence of adverse effects across all investigated endpoints, the limited responses observed in the ECIS assay were primarily linked to elevated concentrations of formulation material. These observations support current regulatory concepts proposing that RNA active ingredients and formulation matrices should be evaluated as distinct assessment entities rather than as a single product characteristic (OECD, 2020; OECD, 2023; Christiaens et al., 2022; De Schutter et al., 2022).

The combined assay strategy presented here provides a practical framework for early-stage hazard identification of RNA-enabled crop protection technologies. By integrating complementary endpoints covering plant stress signaling, formulation biology, and non-target toxicity, this approach enables the detection of unintended biological activity before progressing to higher-tier organismal studies. Such tiered testing strategies may help support future efforts towards harmonized, science-based risk assessment frameworks for RNA-based plant protection products.

## Acknowledgements

M.D., A.K., M.B., A.P. are supported by the Deutsche Forschungsgemeinschaft (DFG, SPP2416, KO 4874/5-1). This work was also supported by the DFG, SPP2416 CodeChi grant (RO 3550/19-1 to S.R. K.E., C.R. and B.M.M. acknowledge financial support by the Deutsche Forschungsgemeinschaft (DFG, German Research Foundation) in the framework of the priority program SPP 2416 “CodeChi” grant MO 476/12-1 (https://codechi.de/). S.F. and J.W. acknowledge financial support by the German Federal Ministry of Education and Research (BMBF) and Bavarian Ministry of Economic Affairs, Regional Development and Energy (StMWi) in the initiative “Biogene Wertschöpfung und Smart Farming" conducted by the Fraunhofer Society. The project was also supported by the High Performance Center Secure Intelligent Systems (LZSiS). We are very much indepted to Prof. Gernot Laengst (Universitaet Regensburg) who provided Sf21 cells used in the assays. J.N., A.C.U.F and A.K.. are supported by Bavarian State Ministry of Food, Agriculture, Forestry, and Tourism (StMELF) and Federal Ministry of Agriculture, Food and Regional Identity.

