## Supplements for "Integrated risk assessment reveals low biological activity of RNA-based crop protection formulations across plant and non-target systems"

### Supplement

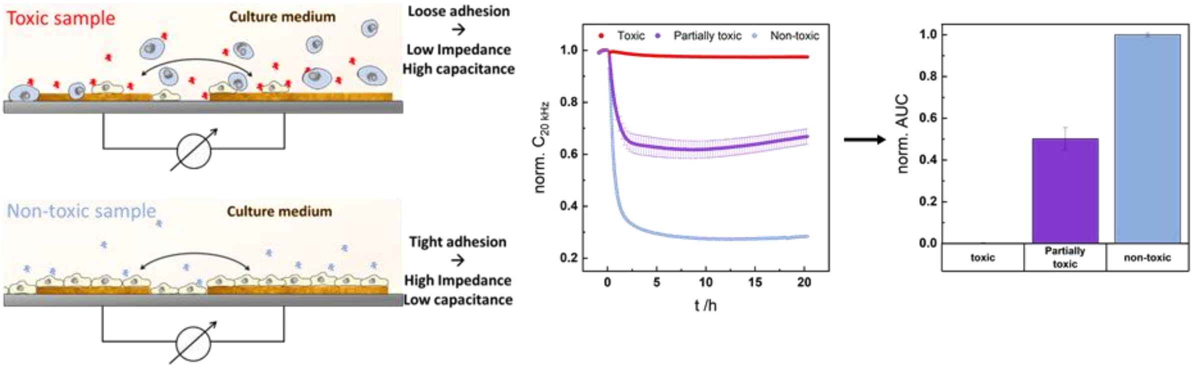


**Supplementary Figure 1: Principle of the ECIS-based phenotypic assay monitoring the time course of cell spreading upon exposure to the compounds under test.** The sketch illustrates raw and processed data for toxic, partially toxic and non-toxic exposures. Time course raw data is analysed by quantifying the area under the curve (AUC).

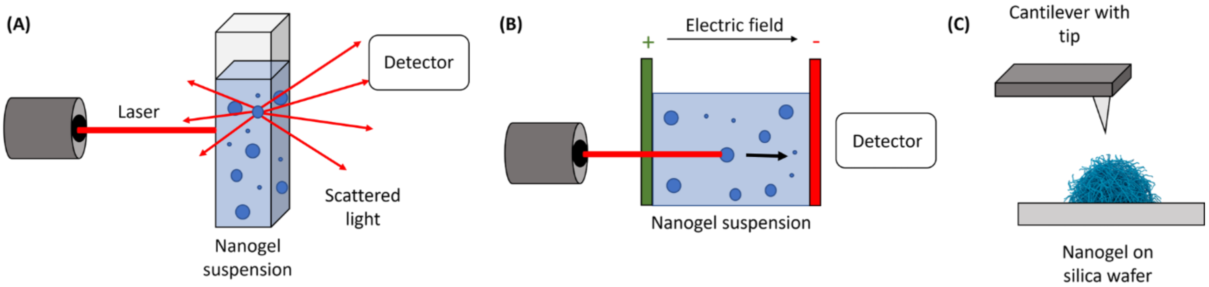


**Supplementary Figure 2: Schematic illustration of the characterization techniques used for chitosan-adipic acid nanogels. (A)** Dynamic light scattering (DLS) measurement of a nanogel suspension to determine the hydrodynamic diameter (Dh) and polydispersity index (PDI). **(B)** Electrophoretic light scattering (ELS) measurement of a nanogel suspension to determine the surface charge, expressed as the zeta potential (ζ). **(C)** Schematic representation of atomic force imaging (AFM) imaging of an individual nanogel deposited on a substrate.

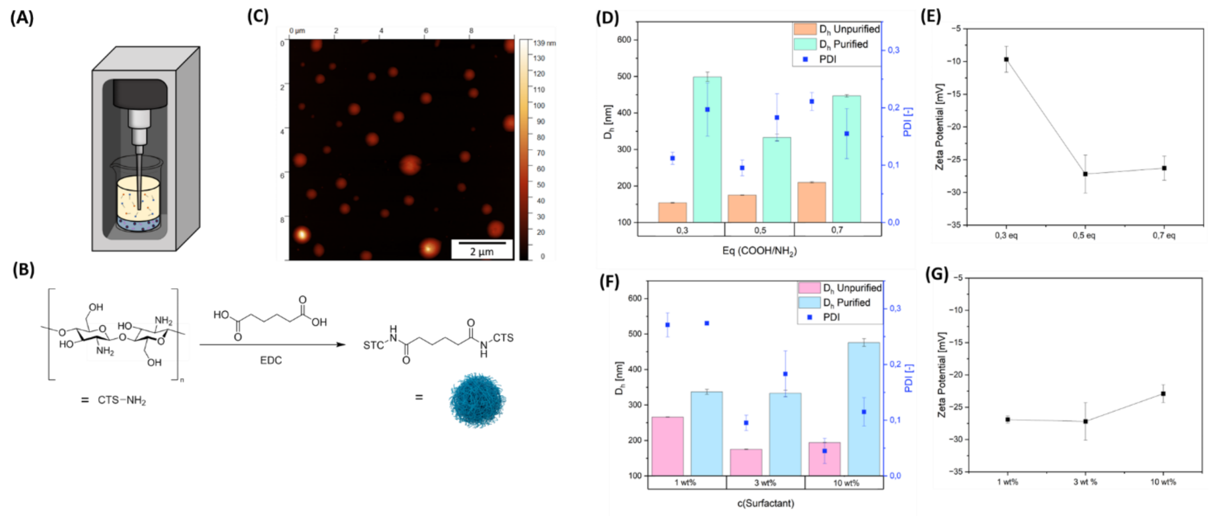


**Supplementary Figure 3: Synthesis and characterization of chitosan-adipic acid nanogels. (A)** Schematic illustration of the inverse miniemulsion synthesis, showing the biphasic reaction mixture before emulsification. **(B)** Schematic representation of the EDC-mediated crosslinking reaction between chitosan and adipic acid. **(C)** Representative dry-state AFM image of chitosan-adipic acid nanogels (MBX-26). **(D)** Average hydrodynamic diameter (Dh) and polydispersity index (PDI) of nanogels prepared with different crosslinking densities ($COOH/NH_2$ = 0.3, 0.5, and 0.7) at a constant surfactant concentration of 3 wt%, measured before purification in the organic phase and after purification in water. **(E)** Corresponding zeta potentials of the nanogels shown in (D). **(F)** Average hydrodynamic diameter (Dh) and polydispersity index (PDI) of nanogels prepared with varying surfactant concentrations (1, 3, and 10 wt%) at a constant crosslinking density ($COOH/NH_2$ = 0.5), measured before purification in the organic phase and after purification in water. **(G)** Corresponding zeta potentials of the nanogels shown in (F). Data are presented as mean ± standard deviation.

**Table 1**: Composition of the aqueous and continuous phases used for the preparation of chitosan-adipic acid nanogels. These formulations were subsequently used for biological characterization and toxicity assessment

| Sample | Eq(COOH/NH2) | Eq(EDC/COOH) | m(CTS) [mg] | m(AA) [mg] | m(EDC) [mg] | c(surfactant) [wt%] |
|---|---|---|---|---|---|---|
| MBX-22-1 | 0.5 | 3 | 20 | 4.41 | 34.7 | 3 |
| MBX-22-2* | 0.5 | 3 | 20 | 4.41 | 34.7 | 3 |
| MBX-23 | 0.5 | 3 | 20 | 4.41 | 34.7 | 10 |
| MBX-24 | 0.5 | 3 | 20 | 4.41 | 34.7 | 1 |
| MBX-25 | 0.3 | 3 | 20 | 2.65 | 20.82 | 3 |
| MBX-26 | 0.7 | 3 | 20 | 6.17 | 48.59 | 3 |

*In contrary to the other samples, MBX-22-2 was added dropwise into 10 mL of deionized water after centrifugation and resuspension in 2 mL ethanol to test stabilizing effects of Tween 80.

**Table 2.** Summary of culture conditions used for the maintenance of Sf21 cells employed in the ECIS-based phenotypic toxicity assays.

| Parameter | Sf21 |
|---|---|
| Culture medium | Sf-900™ II SFM |
| Supplements | None |
| Temperature (°C) | 26 |
| Atmosphere | Dry, no $CO_2$ |
| Cultivation | Suspension, Erlenmeyer flasks |
| Subcultivation | Every 2–3 days |
